# Circulating miRNAs associated with bone mineral density in healthy adult baboons

**DOI:** 10.1101/2021.02.27.433170

**Authors:** Ellen. E. Quillen, Jaydee Foster, Anne Sheldrake, Maggie Stainback, Todd L. Bredbenner

## Abstract

MicroRNAs (miRNAs) regulate gene expression post-transcriptionally and circulate in the blood, making them attractive biomarkers of disease state for tissues like bone that are challenging to interrogate directly. Here we report on five miRNAs – miR-197-3p, miR-320a, miR-320b, miR-331-5p, and miR-423-5p – that are associated with bone mineral density (BMD) in 147 healthy adult baboons. These baboons range in age from 15 to 25 years (45 to 75 human equivalent years) and were 65% female with a broad range of BMDs including a minority of osteopenic individuals. miRNAs were generated via RNA sequencing from buffy coats collected at necropsy and areal BMD evaluated via DXA of the lumbar vertebrae post-mortem. Differential expression analysis controlled for the underlying pedigree structure of these animals to account for genetic variation which may be driving miRNA abundance and BMD values. While many of these miRNAs have been associated with risk of human osteoporosis, this finding is of interest because the cohort represent a model of normal aging and bone metabolism rather than a disease cohort. The replication of miRNA associations with osteoporosis or other bone metabolic disorders in animals with healthy BMD suggests an overlap in normal variation and disease states. We suggest that these miRNAs are involved in the regulation of cellular proliferation, apoptosis, and protein composition in the extracellular matrix throughout life. However, age-related dysregulation of these systems may lead to disease causing associations of the miRNAs among individuals with clinically defined disease.

---

Research interest in microRNAs (miRNAs) has exploded in the past decade with recognition of the potential for these small circulating RNAs to act as biomarkers of disease state or regulators of bone across cell types. As post-transcriptional regulators, these small non-coding RNAs are uniquely able to regulate gene expression in the cytoplasm and may travel among cells to do so. While a number of studies have focused on the potential of miRNAs as biomarkers for osteoporosis and fracture risk, little is known about the miRNAs in the context of normal variation in spinal bone mineral density (BMD) among healthy adults. Here we report on five circulating miRNAs that are associated with variation in BMD within the healthy range for middle-aged and older baboons.

The baboon is unique model for age-related bone loss, as it is closely related phylogenetically to humans, is relatively large bodied, and exhibits intracortical remodeling throughout life, unlike rodent models of skeletal aging. The adult bone remodeling and skeletal fracture properties of baboons are also more similar to humans than are other mid-to large-sized mammals (Wang, Mabrey, and Agrawal 1998; Brommage 2001). Like humans, baboons undergo postmenopausal bone loss and sex and age influence BMD (L Havill et al. 2003) leading to osteopenia in approximately 25% of older females (L. M. Havill et al. 2008). Furthermore, biomechanical properties directly relevant to fracture, including vertebral trabecular bone mechanical properties (LM Havill, Allen, and Bredbenner 2010), femoral cortical bone microstructure (L. Havill et al. 2008), and femoral bone shape (Hansen et al. 2009), are strongly heritable in the baboon, making pedigreed animals ideal for assessing the effects of genetic and non-genetic factors bone biomarkers to bone fragility (Cox et al. 2013).

We leveraged the unique anatomical and genetic features of the baboon to identify circulating miRNAs associated with BMD across a broad range of mildly osteopenic and healthy adult baboons while controlling for underlying genetic factors.

## Methods

### Study Population

We generated miRNA and BMD data for 147 middle-aged and older baboons (hybrid *Papio hamadryas* species) drawn from a larger pedigree of baboons housed at the Southwest National Primate Center and Texas Biomedical Research Institute (Vagtborg 1973). Animals were 65% female and ranged in age from 15 to 25 years of age which is the equivalent of approximately 45 to 75 human years (Figure 1).

**Figure 1.**
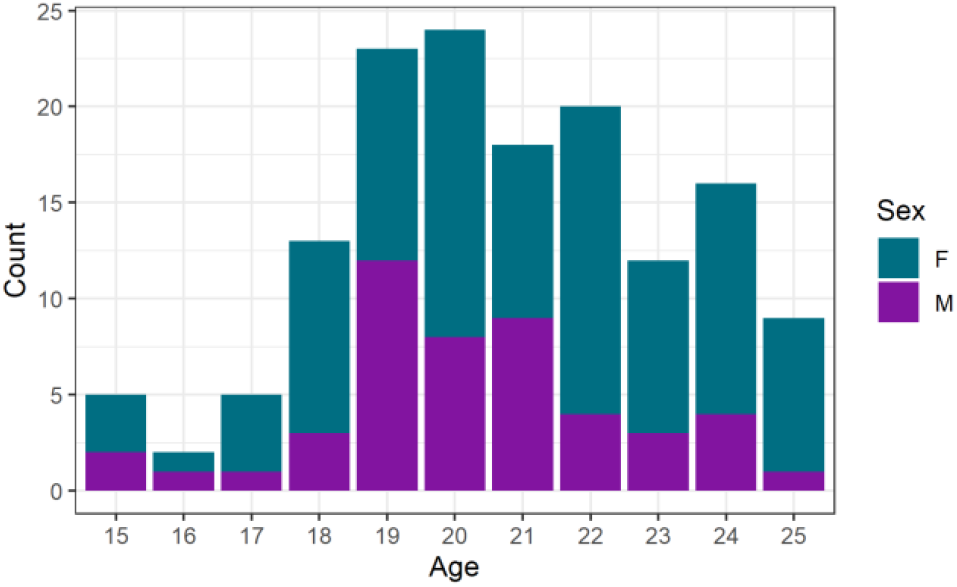
Distribution of animals by age and sex in sample.

During life, all animals were housed outdoors in large social group cages and maintained on commercial monkey chow to which they had ad libitum access. Animals with medical conditions known to influence bone metabolism (e.g. diabetes, chronic renal disease) or a history of traumatic fracture were excluded. No animals were sacrificed for this study, all were euthanized for other reasons and monitored by the Institutional Animal Care and Use Committee. Blood samples were drawn in EDTA tubes at necropsy and processed buffy coat stored at −80 °C until RNA was extracted. Lumbar spines were stored at −20 °C.

### DXA

Dual-energy x-ray absorptiometry (DXA) scans were performed post-mortem on thawed lumbar vertebrae using a Lunar DPX 6529 (General Electric) and areal BMD analyzed with the manufacturer’s software for adults. All analyzed values are for DXA measured for the anterior-posterior (AP) axis of L4-L2 vertebrae. Baboons exhibit high levels of sexual dimorphism in weight and body size leading to significant differences in mean BMD (0.22 g/cm^2^, *p* = 1×10^−6^, Figure 2). The distribution of BMD in these animals is consistent with previously published reference standards from more than 650 baboons (L Havill et al. 2003). Within the 10-year age range of our study, there is no significant decline in BMD. Nevertheless, the distribution of BMD across the sample is broad ranging from 4.3 standard deviations below to 8.4 standard deviations above the healthy, young adult mean for females and −3.5 to 9.3 standard deviation for males. To account for sexual dimorphism in BMD, we performed z-scoring on male and female BMD values separately and combined the scaled values for analysis.

**Figure 2.**
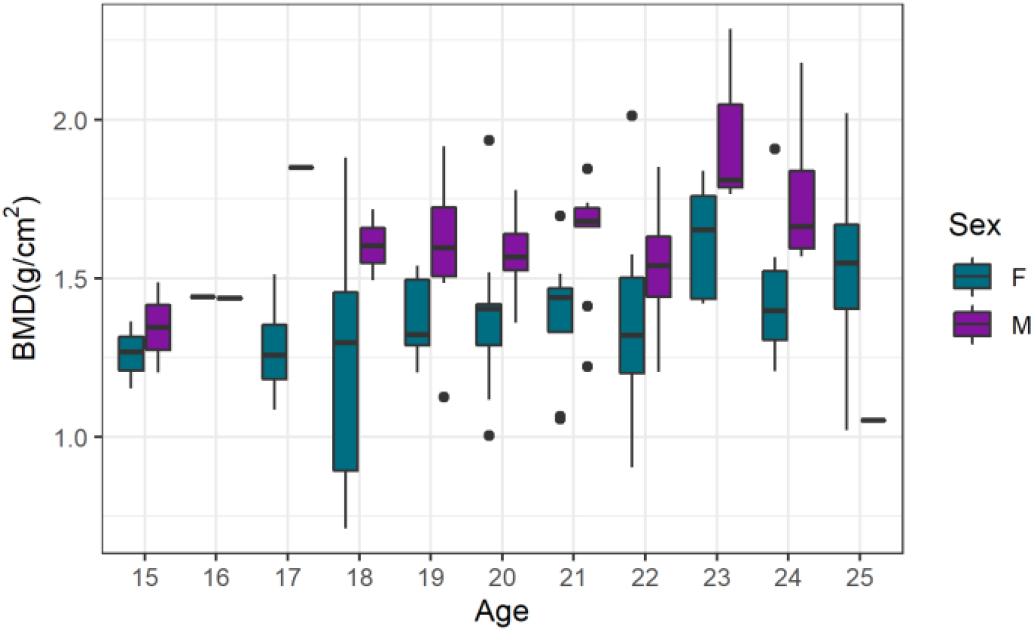
Distribution of BMD in study sample by age and sex.

### miRNA Sequencing and Analysis

Total RNA was isolated from buffy coat using TRIzol (Invitrogen) and the Qiagen miRNeasy Mini Kit as described previously (Spradling et al. 2013). RNA quality was assessed using an Agilent Bioanalyzer 2100 and RNA was enriched for small non-coding RNAs (sncRNAs) by using the Ambion mirVana miRNA Isolation Kit. Complementary DNA (cDNA) libraries were generated with the Illumina Small RNA Prep Kit v1.5 following the manufacturer’s protocol and sequenced in Illumina’s Genome Analyzer (GAIIx) (Karere et al. 2012). mirDeep2 (Mackowiak 2011) was used to align reads to the known human miRbase version 21 (Griffiths-Jones et al. 2008, 2006) mature and hairpin miRNAs and to identify novel miRNAs.

Raw miRNA counts were analyzed using the R statistical package DESeq2 to identify individual miRNAs associated with BMD (Love, Huber, and Anders 2014; R Core Team 2019). DESeq2 has the advantage of incorporating shrinkage estimators for dispersion and fold change which better handles low-abundance miRNAs compared to other methods. After z-scoring, weight and age were no longer associated with BMD and so were not included as covariates. miRNAs significantly associated with BMD at a FDR < 0.05 were further tested to determine if this association was driven by underlying genetic variation in the pedigree. A likelihood ratio test was calculated to compare linear mixed effects models fit with lmekin in the coxme R package (Therneau 2020) with and without miRNA abundance as a fixed effect while including pedigree-based kinship as a random effect.

### miRNA Target Prediction

A major challenge in understanding the potential functional role of miRNAs is that most are computationally predicted to bind dozens of different mRNAs. To better predict what role these circulating miRNAs could play in the bone, we used the R package multiMiR (Ru et al. 2014) to query three experimentally validated miRNA target databases – miRecords (F. Xiao et al. 2009), miRTarBase (Hsu et al. 2011), and TarBase (Karagkouni et al. 2018). To be considered a potential target we required experimental validation of miRNA-mRNA interaction via luciferase, qRT-PCR, or Western blot experiments as well as the presence of the target protein in baboon bone (unpublished mass spectrometry data).

## Results

### miRNAs Associations

After FDR correction, 15 miRNAs were significantly associated with BMD in the DESeq2 analysis of which 5 were associated (likelihood ratio test p < 0.1) after correcting for the genetic relatedness of the animals (Table 1). All five of these highly associated miRNAs miRNAs – miR-197-3p, miR-320a, miR-320b, miR-331-5p, and miR-423-5p – are negatively correlated with BMD (Figure 3) suggesting that increased levels of these miRNAs – which would be expected to decrease mRNA expression – are indicative of a decline in BMD. The heat map in Figure 4 illustrates a high level of heterogeneity in miRNA abundance across animals with varying BMD.

**Table 2.** miRNAs significantly associated with BMD after FDR correction in DESeq2 analysis. Bolded miRNAs remained significantly associated after correction for pedigree structure.

| | $\beta$ | Mean Expression | $p$ | $LRT\ p$ |
| --- | --- | --- | --- | --- |
| **hsa-miR-423-5p** | **-0.28** | **886** | **$2.4 \times 10^{-4}$** | **0.03** |
| **hsa-miR-320a** | **-0.22** | **2,561** | **$3.9 \times 10^{-3}$** | **0.04** |
| hsa-miR-339-5p | -0.19 | 1,697 | $6.2 \times 10^{-3}$ | 0.22 |
| hsa-miR-3173-5p | -0.38 | 16 | $3.0 \times 10^{-2}$ | 0.14 |
| hsa-miR-371a-5p | 0.6 | 7 | $3.0 \times 10^{-2}$ | 0.40 |
| hsa-miR-16-2-3p | -0.44 | 137 | $3.3 \times 10^{-2}$ | 0.27 |
| **hsa-miR-197-3p** | **-0.18** | **577** | **$3.5 \times 10^{-2}$** | **0.09** |
| hsa-miR-21-5p | 0.16 | 30,037 | $3.5 \times 10^{-2}$ | 0.68 |
| **hsa-miR-331-5p** | **-0.09** | **808** | **$3.5 \times 10^{-2}$** | **0.07** |
| hsa-miR-374a-3p | 0.26 | 231 | $3.5 \times 10^{-2}$ | 0.88 |
| hsa-miR-374a-5p | 0.19 | 4,255 | $3.5 \times 10^{-2}$ | 0.92 |
| hsa-miR-424-3p | -0.16 | 86 | $3.5 \times 10^{-2}$ | 0.10 |
| hsa-miR-484 | -0.13 | 2,334 | $3.5 \times 10^{-2}$ | 0.19 |
| **hsa-miR-320b** | **-0.21** | **28** | **$3.6 \times 10^{-2}$** | **0.05** |
| hsa-miR-574-3p | -0.16 | 94 | $4.1 \times 10^{-2}$ | 0.25 |

**Figure 3.**
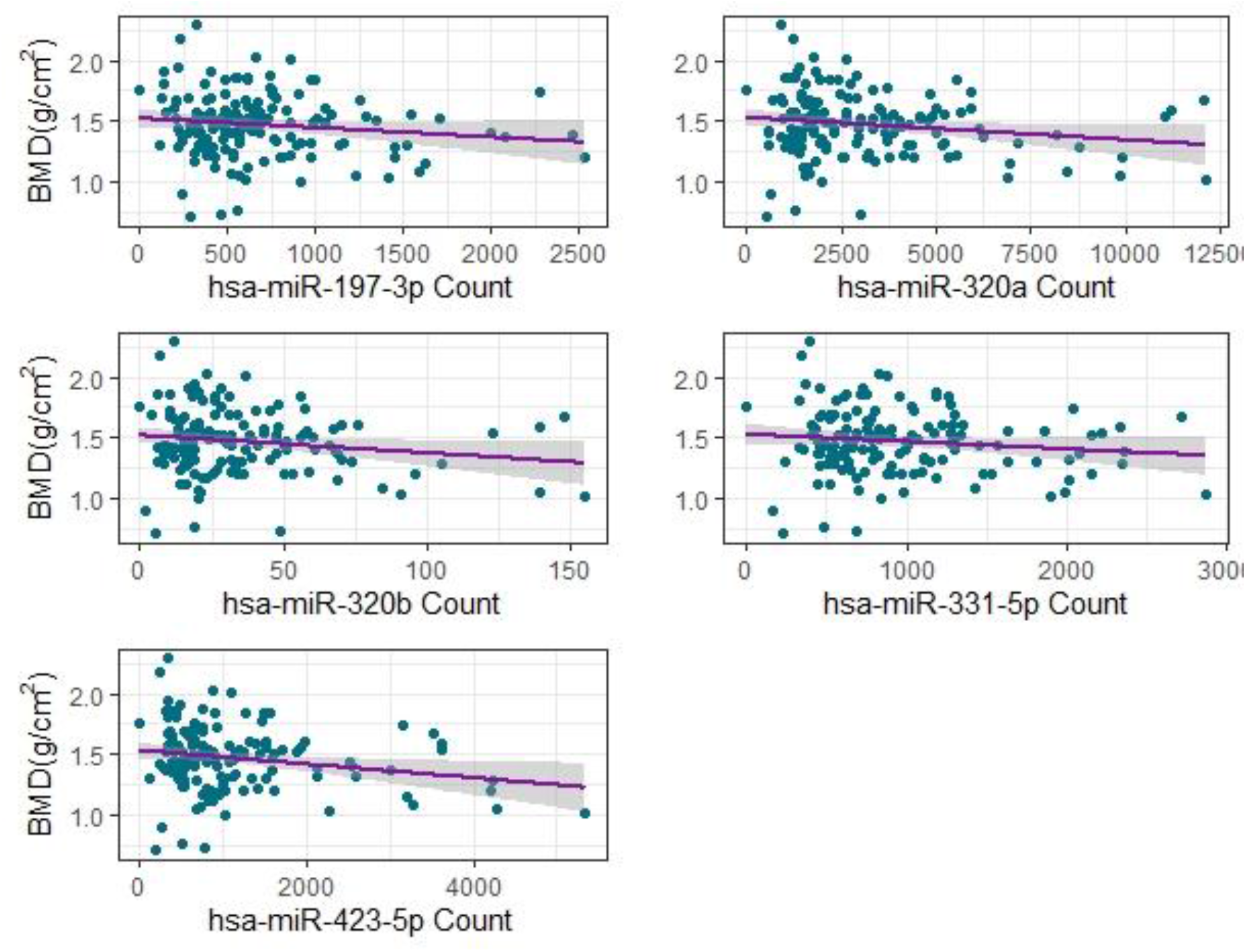
miRNA abundance versus BMD for significantly associated miRNAs.

**Figure 4.**
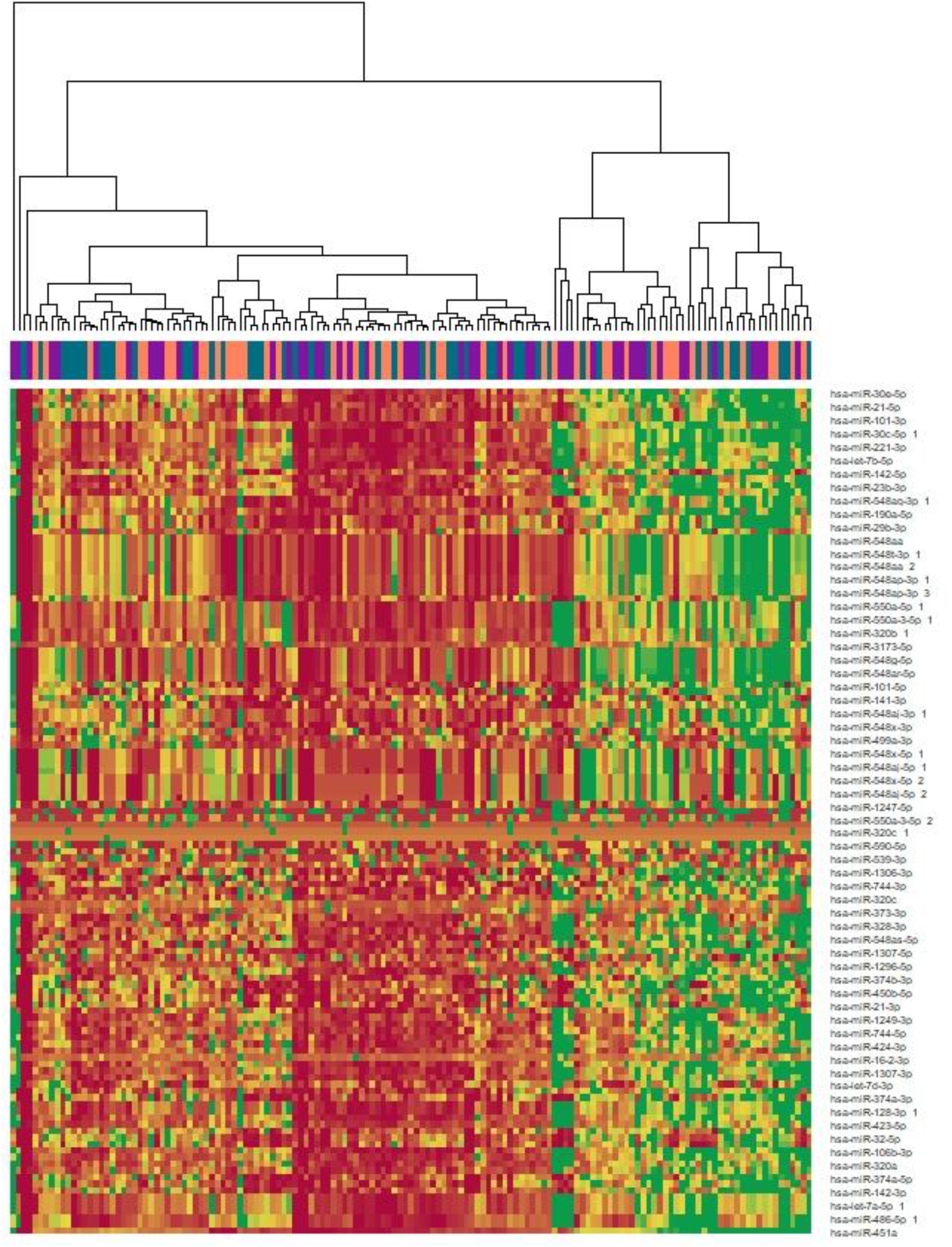
Heatmap of all miRNAs with nominal *p* < 0.05 in DeSeq2. Columns are color-coded by z-scored BMD tertile with purple indicating the lowest tertile and teal the highest.

### Putative miRNA targets

Based on our screening criteria, we identified 60 genes present in bone and experimentally validated as potential targets of our five significant miRNAs (Supplementary Table 1). Analysis in DAVID (Huang, Sherman, and Lempicki 2009) identified several significantly enriched functional annotations within the 60 genes. The top five enriched gene ontology terms representing more than 10% of the genes on the list are myelin sheath (26.4 fold enrichment, FDR = 6.1×10^−10^), focal adhesion (11.2 fold enrichment, FDR = 2.9×10^−7^), extracellular matrix (11.1 fold enrichment, FDR = 3.6×10^−5^), cadherin binding involved in cell-cell adhesion (9.5 fold enrichment, FDR = 9.0×10^−4^), cell-cell adherens junction (9.0 fold enrichment, FDR = 5.4×10^−4^). Additionally, these putative targets include seven genes previously linked to osteoporosis in genome wide association studies based on data drawn from the Public Health Genomics and Precision Health Knowledge Base (v7.1) (Yu et al. 2010) (Table 2).

**Table 2.**
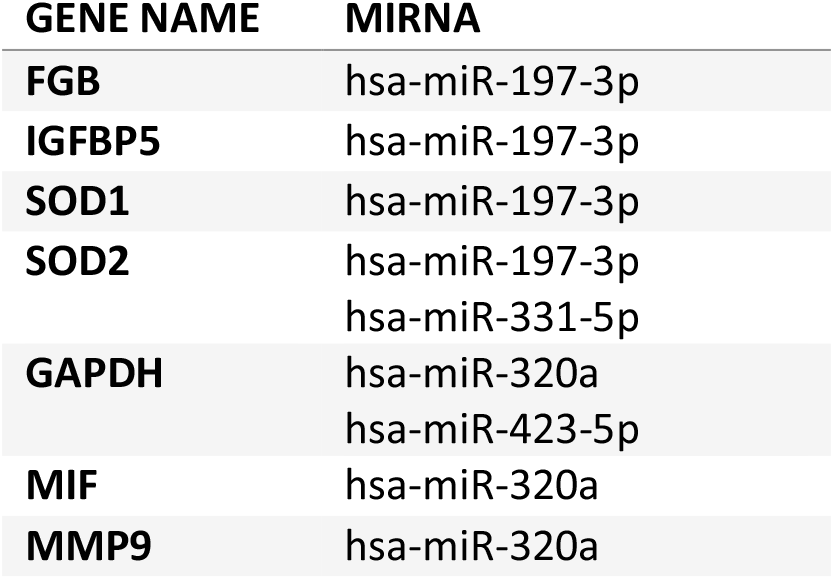
miRNA targets linked to osteoporosis in genome wide association studies

| GENE NAME | MIRNA |
| --- | --- |
| **FGB** | hsa-miR-197-3p |
| **IGFBP5** | hsa-miR-197-3p |
| **SOD1** | hsa-miR-197-3p |
| **SOD2** | hsa-miR-197-3p<br>hsa-miR-331-5p |
| **GAPDH** | hsa-miR-320a<br>hsa-miR-423-5p |
| **MIF** | hsa-miR-320a |
| **MMP9** | hsa-miR-320a |

## Discussion

Most previous research on bone density and miRNAs has focused on patients with osteoporosis or other disorders of bone metabolism. This is reflected in links between osteoporosis or fracture risk in human patients and the miRNAs we have identified, many of which have been suggested as potential clinical biomarkers.

Of the miRNAs we identified, literature linking the miRNA-320 family to bone fragility is the most robust. miR-320b is more common in women with a recent fracture compared to age-matched controls (Weilner et al. 2015). This may be due to its role in osteoblast differentiation. miR-320b over-expression prevents osteoblast differentiation, while inhibition promotes bone matrix mineralization and differentiation via BMP-2 (Laxman et al., 2017). Additionally, researchers studying low-traumatic fractures in premenopausal, postmenopausal, and idiopathic osteoporosis patients found miR-320a was upregulated in these patients compared to controls (Kocijan et al., 2016). Prior analysis of trabecular bone tissue showed significant dysregulation of miR-320a in patients with osteoporosis, possibly via regulation of osteogenesis by targeting bone-forming genes such as *CTNNB1* (B-catenin) and *RUNX2* (De-Ugarte et al., 2015). These findings led to the inclusion of miR-320a on the commercially available OsteomiR panel (Tamirna).

Similarly, in a study of plasma miRNAs associated with fracture risk, miR-423-5p expression was significantly negatively associated with FRAX score, but not BMD, in osteoporosis patients (Bedene et al. 2016). This finding was reinforced by research on facial bone atrophy where miR-148-3p was shown to promote bone proliferation via prevention of apoptosis in mesenchymal stem cells (Yang et al. 2018). While there is little additional evidence of a role for this miRNA in bone, it has also been linked to regulation of apoptosis in cardiomyocytes (Zhu and Lu 2019), kidney cells (Yuan et al. 2017), retinal pigment epithelial cells (Q. Xiao et al. 2019), and colon cancer cells (Jia et al. 2018).

Circulating miR-331-5p levels have been identified as a potential biomarker for osteoporosis and subsequent bone fracture (Liu et al. 2015), although there is little known about the role of this miRNA in bone. Hints to its function come from studies of vascular smooth muscle cells, where miR-331-5p is induced by BMP2-PPARγ signaling to regulate cellular proliferation (Calvier et al. 2017). We predict a role for miR-331-5p in regulating *SOD2* expression which directly induces a BMP2 response under hypoxic conditions (Kamiya et al. 2013).

While miR-197-3p has not previously been linked to adult bone metabolism, it was identified in the downregulation of osteogenesis in human amniotic membrane-derived mesenchymal stem cells due to its suppression of SMAD2 in the TGF-β pathway during osteoblast differentiation (Avendaño-Félix et al. 2019). We predict that miR-197-3p may target expression of both *SOD1* and *SOD2*, antioxidative enzymes that respond to vitamin D levels (Lisse 2020) and are thought to play a critical role in the regulation of cellular senescence (Zhang et al. 2017; Jeong and Cho 2015).

Taken together, our BMD-associated miRNAs and their putative targets point towards regulation of extracellular matrix proteins, apoptosis, and cell proliferation – three key components in the maintenance of bone homeostasis. Our findings demonstrate the overlap in miRNAs associated with bone mineral density and related traits in humans and nonhuman primates, highlighting the utility of this model for understanding aging bone. The association of these miRNAs with variation in bone mineral density within the healthy, non-osteoporotic, range suggests they may be useful in identifying the earliest stages of metabolic shifts in bone. While the impact of epigenetic regulation of gene expression is evident in bone tissue, the use of miRNAs as biomarkers for low BMD and high fracture risk is still relatively new. It is apparent more research is needed to better understand these molecular pathways. Future work should focus on identifying the presence and functional role of these miRNAs in bone tissue to solidify their promise as biomarkers.

## Acknowledgements

We thank Lorena Havill for her role in the development of this project. This work was funded by R01 AR064244 to TLB and K01 AG056663 to EEQ. The animals in the SNPRC pedigree were supported by grant P51 OD011133.

